# Sleep patterns predicting stress resilience are dependent on sex

**DOI:** 10.1101/2024.12.30.630808

**Authors:** Brittany J Bush, Affra Mohamed, Eva-Jeneé Andrews, Gabrielle Cain, Ayobami Fawole, Hadiya Johnson, Ashton Arocho, Zhimei Qiao, Ketema N Paul, J. Christopher Ehlen

## Abstract

Sleep disturbances and stress have a well-established link with neuropsychiatric illness; however, the nature of this relationship remains unclear. Recently, studies using the mouse social-defeat stress model revealed a causal role for non-rapid eye movement (NREM) sleep in the maladaptive behavioral responses to stress. These results suggest a novel function for NREM sleep; as a response by cortical neurons to mitigate the maladaptive effects of stress. A major limitation in many social defeat studies has been the exclusion of females. Women exhibit a greater prevalence of both affective disorders and sleep disturbances compared to men, thus there is a clear need to understand sleep – stress interactions in females. The present study adapts recently developed female social-defeat stress models to allow social-defeat and EEG in male – female pairs. Our findings duplicate the behavioral responses that occur in other female, nondiscriminatory, and male models of social-defeat stress. Analysis of electroencephalographic (EEG) recordings, before exposure to stress, reveal that resilience is associated with differences in both NREM and REM sleep that are dependent on sex. After social defeat stress, NREM sleep was increased only in resilient males. In females, susceptibility to stress was associated with increased durations in NREM-sleep bouts. A potential cause of these sleep differences was also identified prior to stress exposure, sex differences in recovery from NREM-sleep loss; thus, suggesting an underlying sex-difference in the homeostatic process regulating sleep interactions with social-defeat stress. These findings suggest that NREM sleep quality is lower in resilient females, whereas the amount of REM sleep is decreased in susceptible females—when compared to males of the same behavioral phenotype. Overall, our findings reveal sexual dimorphism in both the sleep characteristics predicting resilience and sleep changes induced by social-defeat stress.

## Introduction

Both sleep disturbances and stress have a well-established link with neuropsychiatric illness, but the causal nature of this relationship is not clear. The social-defeat stress model is frequently employed to investigate core features of stress-induced neuropsychiatric illnesses. This model uses a resident-intruder paradigm wherein a mouse (intruder) experiences social-defeat stress in the home cage of a larger mouse (resident)[1]. Social-defeat stress leads to a sustained syndrome of maladaptive behaviors; including anhedonia, increase in anxiety-like behavior, metabolic alterations [2], changes in circadian rhythms [3] and, of importance to this application, the avoidance of social interaction [1, 4]. Notably, there are subpopulations of mice that are naturally resilient to the development of these maladaptive behaviors, thus this model allows investigations into the mechanistic origins of stress-induced behavior.

The social-defeat stress model has significantly advanced our understanding of maladaptive behavioral responses to stress, but most of this research focuses on males. Furthermore, an overall lack of data in females is a major gap in our mechanistic understanding of neuropsychiatric illness. It is essential to understand the interaction of sleep and stress in females as women exhibit a two-fold greater prevalence of affective disorder [5–8], increased occurrence of insomnias, and report poor sleep quality more frequently than men [9–13]. Furthermore, women diagnosed with an affective disorder report problems with sleep quality at a greater rate than men [14–16]. These finding illustrate a clear need to investigate sex-differences in maladaptive behavioral responses to stress.

Sleep responses to non-social stress modalities have provided insights into the reciprocal relationships between stress and sleep responsivity. In rodent models, exposure to non-social stress modalities have a variety of effects on non-rapid eye movement (NREM) and rapid eye movement (REM) sleep [17]. The specific effects on sleep are dependent on the type of stressor (e.g., immobilization, electric shock, open field, etc.) and other factors, such as the frequency and duration of stress [18]. The first study of the effects of stress on sleep in male rats reported that two hours of immobilization stress increased REM sleep and total sleep, primarily during the active (wake) phase of the sleep-wake cycle [19]. A subsequent study in mice recapitulated these effects in male mice but also revealed that female mice did not exhibit a similar increase in REM sleep and total sleep [20]. This study underscores the limitations of studies that do not include females and do not examine sex differences in sleep responses to stress. More recently, total sleep deprivation has been demonstrated to reduce fear responses to non-social stress modalities in male and female mice [21], providing additional evidence that sleep amount prior to stress exposure has regulatory influences on stress responsivity.

A bidirectional relationship exists between social-defeat stress and sleep. Social stress causes increased NREM sleep and NREM sleep intensity (slow-wave activity [SWA]; [22, 23]. Furthermore, NREM sleep measured by electroencephalography (EEG), prior to stress exposure, predicts resilience to social-defeat stress; these predictive factors include NREM fragmentation, NREM sleep intensity, and local electrical activity in the medial prefrontal cortex (mPFC) [24–27]. A causal relationship was also demonstrated between sleep and resilience; experimentally increasing or decreasing NREM sleep duration (during the natural sleep period) promotes or inhibits respectively, resilience to social defeat stress [27]. Overall, these results reveal a novel function of NREM sleep; as an active response by cortical neurons to mitigate the effects of stress, however, the mouse model used in these studies did not allow investigations in females. In the present study we adapted a newly available female model of social-defeat stress to incorporate EEG recordings. Notably, this model allows simultaneous social-defeat and EEG in both females and males [28]. Using this model, we evaluate pre-existing sleep differences that predict resilience to social defeat stress and measure sleep changes occurring immediately following defeat stress. The results provide a direct comparison of sleep between males and females with known responsiveness to stress (resilient or susceptible to social-defeat stress) both before and after exposure to social-defeat stress.

## Methods

### Animals

Male and female C57/BL6J mice purchased from Jackson Laboratories (Bar Harbor, ME) were between the ages of 7 to 9 weeks at the start of the study, weighing approximately 18-25g. Male, CD-1 mice were used as aggressor mice during the social defeat stress paradigm and were purchased from Charles Rivers Laboratories (Wilmington, MA). CD-1 mice were retired breeders and greater than three months in age, weighing approximately 30-35g. All mice were maintained on a 12:12 L:D light cycle and had food and water available *ad libitum*. All C57/BL6J mice were housed as male – female pairs, in clear plastic cages, separated by a perforated barrier, throughout the study (social avoidance testing was conducted individually). All procedures involving animals received prior approval from the Morehouse School of Medicine Institutional Animal Care and Use Committee.

### Electroencephalogram (EEG) Surgery

Mice were anesthetized with isoflurane and the head of the mouse was secured during EEG surgery using a stereotaxic frame. Prefabricated 3-lead EEG implants, purchased from Pinnacle Technologies, Inc (Lawrence, KS), were anchored to the skull with four stainless steel screw (0.10 – 0.12mm long) that also served as epidural electrodes. Pairs of screw-electrodes were placed bilaterally approximately 1.5mm anterior or 2.5mm posterior to bregma. Electromyographic (EMG) electrodes were inserted into the nuchal muscle for electromyographic recordings. Continuity between the screw-electrodes and implant was maintained by silver epoxy. Dental acrylic was used to cover the implant and the wound closed with nylon suture. Subcutaneous carprofen was used for analgesia (daily for three-days).

### Sleep Restriction/Sleep Deprivation

After two weeks of recovery from surgery, one 24 – hour undisturbed baseline EEG recording was obtained and followed by six hours of sleep restriction from Zeitgeber time (ZT) 0-6 (ZT0 = lights on). Wakefulness was maintained in both male and female mice by gentle handling, and the introduction of novel objects into the cages by a trained observer. The amount of sleep lost was calculated as the difference between sleep amount during the six-hour sleep deprivation and sleep amount in undisturbed conditions (measured during the same time) on the previous day.

### Female Social Defeat Stress Model and Aggressor Training

Social defeat stress occurred for ten consecutive days; consisting of three daily defeats (five-minute duration) separated by five–minute breaks. Each male – female pair was placed into the home cage of a novel, aggression-tested, male CD-1 mouse (aggressor) for each five-minute defeat. Defeats were monitored continuously, and mice were separated for 10 seconds if excessive aggression (>15 seconds of consistent contact) occurred. Wounding was rare, if small bite wounds were found, the defeat session was ended and betadine applied to the affected area. Three males received one bite wound during defeats, one of these males was from the group of five that did not undergo sleep recording. All defeat procedures were performed during the first two hours of the dark period (ZT12-14). Dim red light (<5 lux) was used for procedures conducted in the dark. This social defeat stress procedure was modified from the previously published procedure of Yohn, et al., 2019 (N.b. mice were not continuously housed with an aggressor between social-defeat stress sessions and three, 5 – minute defeats were used instead of one 5 – minute defeat). Within the week prior to social defeat stress, CD-1 mice were tested for aggression using a five-minute interaction with a male – female, C57/BL6J, mouse pair for three consecutive days. CD-1 mice that displayed aggression for at least two of the three interactions (latency to display aggression < 1 min) were used in the study.

### Social Interaction Testing

Social interaction testing was conducted in a 30×30 cm arena that held a caged, novel, CD-1 mouse (9×9cm cage). Testing was performed on the first day of the study and again two days after the last session of social defeat stress (days 1 and 18; **Figure 1A**). C57/BL6J mice were placed into the center of the arena for three minutes in consecutive trials with and without the CD-1 mouse present. The positioning of the C57BL/6J mouse during the social interaction test was monitored and the time spent in standardized regions of the arena was used to (Noldus Ethovision XT) calculate a social interaction ratio as follows:

**Figure 1.**
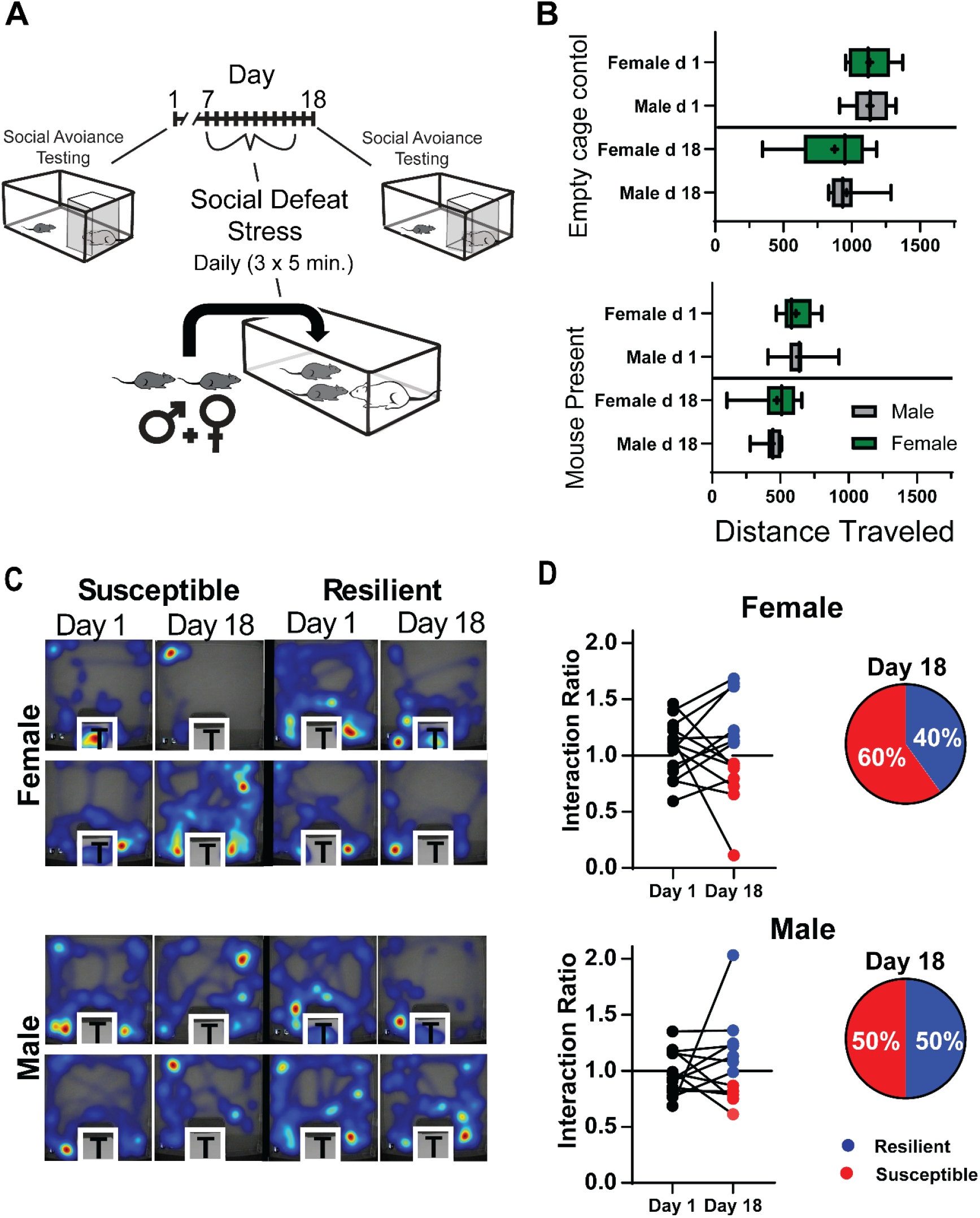
Social interaction testing pre and post social-defeat stress: Study paradigm showing the timing of social interaction testing both before and two days after the ten days of social defeat stress (A). Seep recordings and sleep restriction were also performed but are not included in this figure’s timeline; see figure two. Distance traveled in the testing arena was not different between males and females during any of these tests (B). This was true for testing without (top, empty cage) and with (bottom, mouse present) a caged novel CD-1 mouse in the testing arena. Representative heatmaps of individual tests with a CD-1 mouse present (C; warmer colors represent more time). Social interaction were ratios calculated as time spent near the CD-1 mouse during the three – minute test divided by time spent near the empty cage (D). Interaction scores greater than 1.1 indicated resilience, less than 0.9 indicated susceptibility. Box plot (B) shows interquartile range, mean (+), median (line) and min-max. n=10 males, n=14 females. Additional analysis can be found in Figure S1. Statistical details can be found in Table S1.

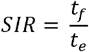

Where *t*_*e*_ = time within 15 cm of the empty cage, and *t*_*f*_ = time within 15 cm of the caged, novel CD-1 mouse. A ratio >1.1 confirmed resilience, while a ratio <0.9 confirmed susceptibility. Mice with interaction ratios between 0.9 and 1.1 after social defeat were excluded from sleep analyses but are included in behavioral analysis (n = 1 female, n = 1 male).

### Sleep Acquisition and Scoring

Seven days following EEG surgery, mice were transferred to housing as male – female pairs in perforated, open top sleep recording cages. Each C57/BL6J mouse in the pair was individually connected to a lightweight recording tether and low resistance commutator (Pinnacle Technologies, Inc.) secured above the open top cages. This allowed complete freedom of movement around the cage. After seven days of acclimation to the sleep recording system, 24 – hours of baseline EEG/EMG data were recorded (Acquisition software, Pinnacle Technologies, Inc.). EEG/EMG signals (EEG low-pass filtered; 30 Hz cutoff) were continuously recorded throughout the duration of the study. EEG/EMG recordings were analyzed by a trained observer in ten second epochs and classified as awake, NREM sleep, REM sleep, or artifact based on the waveforms that predominated in each epoch. Wake: low-voltage, high-frequency EEG; high-amplitude EMG; NREM: high-voltage, mixed-frequency EEG; low-amplitude EMG REM: low-voltage EEG with a predominance of theta activity [6–10 Hz]; very low amplitude EMG. Analysis of EEG recordings was conducted by individuals blinded to the experimental treatments and behavioral phenotypes. EEG files with more than 5% artifact were excluded from all analyses (n=1, see statistical analysis section for details). Sleep classification was performed by two scorers who achieved at least 95% agreement on a set of standard training files maintained in our lab and five files chosen randomly from the current data set.

Power spectral analysis was accomplished by applying a fast Fourier transform (FFT, 0.1 Hz frequency resolution) to the raw EEG signal. Normalized slow-wave activity was calculated using spectral power within the 0.5-4Hz (delta) frequency band divided by the 24 – hour average in this same frequency band on the first day of EEG recording (obtained in undisturbed conditions). Relative power expressed 0.5-4Hz power as a percentage of total power in the same recording (0.5-30 Hz) calculated in ten-second epochs. Slow-wave energy (*energy)* measures the accumulation of slow wave activity over time and was calculated using the 0.5-4Hz frequency range as follows:

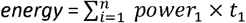

### Statistical Analysis

Fifteen male – female pairs completed the experimental paradigm. EEG recordings were performed on 15 female and 10 male mice. Five male mice used in the study did not undergo EEG surgery. One male and three female mice were excluded from EEG analysis due to technical issues that resulted in > 5% artifact or signal loss during the study; however, all mice that completed the behavioral paradigm were included in the behavioral analysis. EEG data was analyzed using one-way or two-way analysis of variance (ANOVA) with repeated measures as appropriate. Student’s t-test was used for the comparison of two groups. Post-hoc analysis was conducted using the Holm-Sidak method which adjusts α to maintain the family-wise error-rate at 0.05, thus adjusting for repeated comparisons. Significance was defined, in all analysis, as a p-value < 0.05. A Sharpiro–Wilk normality test was used to confirm that data were normally distributed. Statistical results are presented in **Table S1**. An appropriate sample size of six was calculated using a type I error rate of 0.05 and type II error rate of 0.2. This analysis was based on total sleep data (total sleep increase after social defeat stress for susceptible mice) reported in our previous study [22]. The values for standard deviation and mean difference were 14.6 min and 25 min, respectively.

## Results

Males and females experienced aggressive encounters during social defeat stress; however, males were attacked more frequently than females (**Figure S1**). These findings were similar to Yohn, et al., 2019 [23]. Male – female pairs were individually tested for social interaction in an arena containing a caged, novel, CD-1 mouse. This testing was done twice; prior to social defeat stress and two days after the ten-day testing paradigm (**Figure 1A**). Results of this testing are shown in **Figure 1B-C and S1**. No sex differences were found in exploratory activity during testing (measured by total distance traveled in the arena; **Figure 1B**) with or without the presence of a novel mouse. This verifies that general changes in locomotor activity did not influence the social-interaction test. Neither did social interaction scores differ between males and females on either day of testing (Mann-Whitney U: Day 1, U = 69.5, p = 0.47; day 18, U = 83.5, p = 0.99). Social interaction scores determined two days after the last day of social defeat stress (day 18; Figure 1A and 2B; Last social defeat, day 16; undisturbed sleep recordings, day 17) were used to identify mice as susceptible or resilient in the subsequent sleep analysis. Significant behavioral changes were induced by stress in both male and female mice. Our results show increased time interacting with the target mouse when resilient and decreased time interacting when susceptible (**Figure S1**). Furthermore, when interaction ratios of susceptible and resilient mice were examined separately there was a clear pattern of change from pre-to post-social defeat stress, with increased interaction ratios in resilient mice and decreased interaction ratios in susceptible mice (**Figure S1**). An equal number of susceptible and resilient males were identified by social interaction score, whereas more susceptible female mice (60%) were identified after social defeat stress (**Figure 1C, D**).

We first compared baseline sleep between males and females (**Figure 2A, top**), prior to social-defeat stress (**Figure 2B**). Three-way ANOVA was used to compare the main effect of sex, resilience, and time on NREM and REM sleep and slow-wave activity (SWA). The results revealed a significant main effect of sex and time (details in **Table S1**) for both REM sleep and SWA. This led us to collapse the data across the resilience phenotype for further examination (**Figure 2**). NREM sleep amount was not different between males and females; however, males did show elevated NREM slow-wave activity (SWA), during the late active period, when compared to females. We then used social interaction scores (measured at the end of the study, following social defeat stress) to assign mice to susceptible or resilient subpopulations and examined sex-differences in sleep within each subpopulation (**Figure 2B**). Mice that were resilient following social-defeat stress had pre-defeat NREM SWA that was significantly higher in males when compared to females (**Figure 2A, middle**). Mice that were susceptible to social-defeat stress, in contrast, had pre-defeat REM sleep that was significantly higher in males when compared to females (**Figure 2A, bottom**). This increased REM sleep in males was primarily during the sleep (light) period of the 24-hour day.

**Figure 2.**
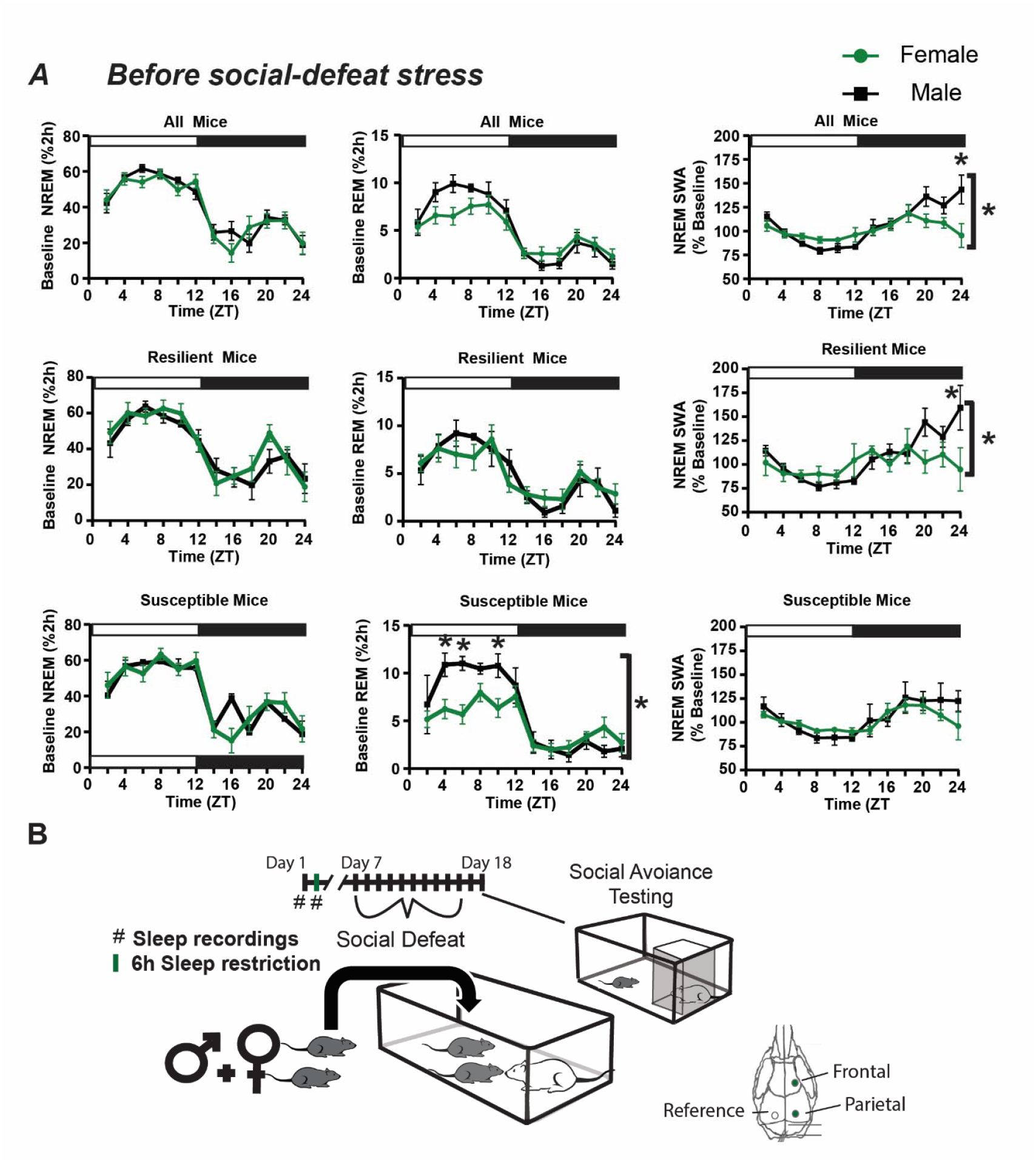
Sex differences in slow wave activity and REM sleep amount before social-defeat stress. The patterns and amounts of non-rapid eye movement (NREM) and rapid eye movement (REM) sleep are shown with NREM slow-wave activity (SWA; A) in undisturbed conditions. An overview of the experimental paradigm is illustrated in B along with the timing and lead arrangement for electroencephalographic (EEG) recording. Slow-wave activity (SWA) during NREM sleep was significantly increased in male mice, compared to females (A, top row). Prior to social-defeat stress (B, day 1), male mice that were resilient to stress (A, middle row) had significantly increased NREM SWA, compared to females (frontal lead shown). This difference was not seen in mice identified as susceptible (A, bottom row). Rapid eye movement (REM) sleep prior to social-defeat stress was significantly greater in male mice that were identified as susceptible after social-defeat stress, when compared to females (A, bottom row). Mice were identified as resilient or susceptible by social interaction testing two days after the ten days of social-defeat stress (B). All plots show mean +/-standard error of the mean (SEM). n=12 females, n=8 males. Side brackets indicate significant ANOVA main effect of sex or interaction p < 0.05; *, p < 0.05, Holm-Sídak’s multiple comparisons test. Statistical details can be found in Table S1.

Pre-defeat sleep was also compared between susceptible and resilient subpopulations, to identify sleep differences associated with resilience (**Figure 3**). No differences in baseline NREM, REM or total sleep amount were found between resilient and susceptible mice, although there was a trend indicating increased REM sleep in susceptible males (**Figure 3A**), but not females. Increased relative NREM SWA was found in resilient males when compared to susceptible males (**Figure 3B**). This increased NREM SWA, a measure of sleep intensity, was only detected in EEG leads placed over the parietal region of the brain. Analysis of sleep bouts also revealed significantly increased REM bout duration in susceptible mice when compared to resilient mice. This difference was significant in the 12h light period of both males and females (**Figure 3C**). Additional sex differences were found in this sleep-bout analysis with males having significantly more NREM bouts than females. Furthermore, total REM bout duration was significantly longer in female mice (**Figure 3C**).

**Figure 3.**
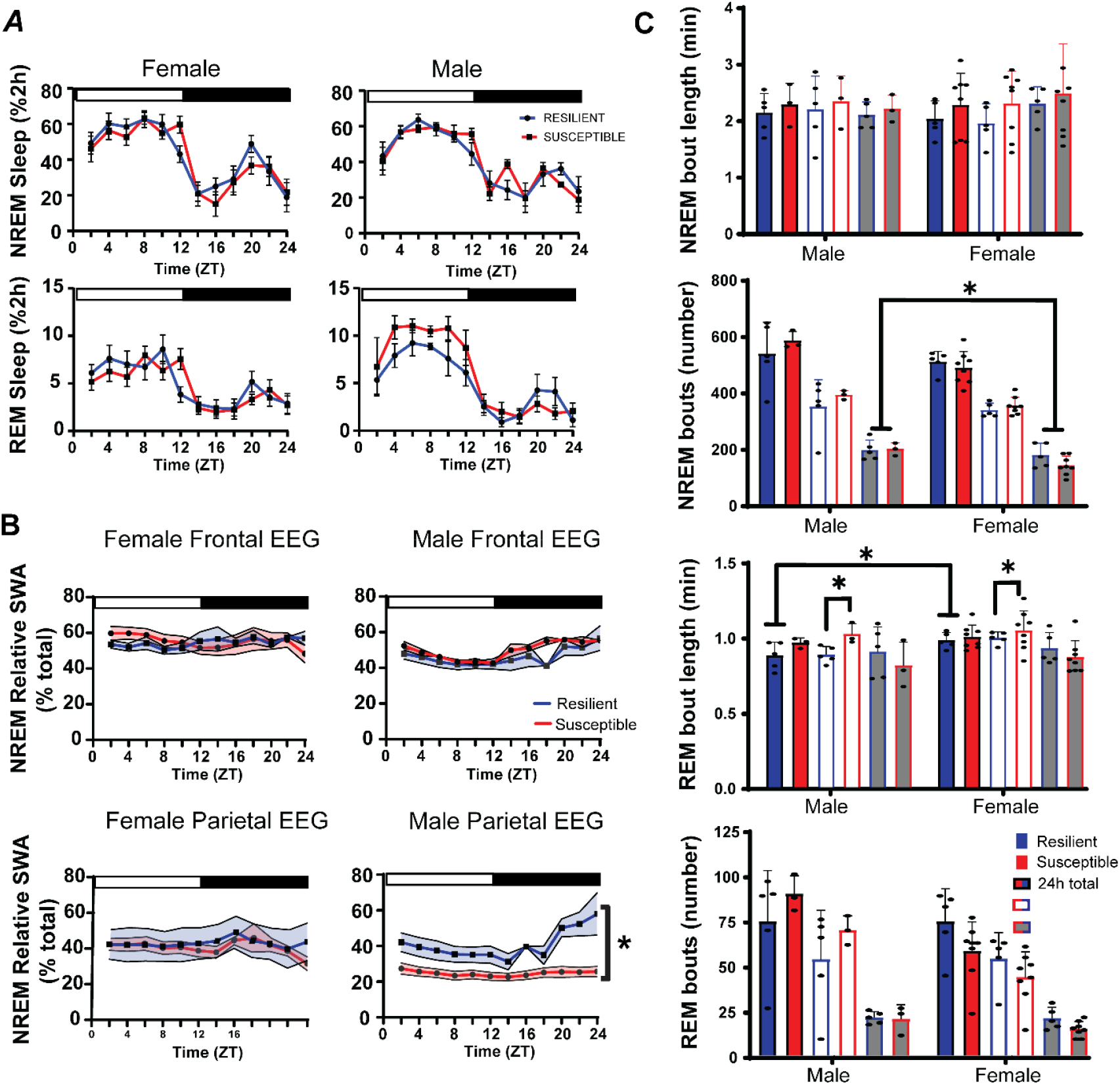
Sleep before social-defeat stress in susceptible and resilient mice. No significant differences in REM or NREM sleep amount we found between susceptible and resilient mice before exposure to social-defeat stress (A); however, NREM relative slow-wave activity (SWA) was significantly greater in resilient male mice when compared to susceptible males. These differences were only found in the parietal EEG lead of males and were not found in females (B). Analysis of sleep bout number and duration (C) revealed significantly more NREM bouts in males during the dark period, compared to females. Sex differences were also found in REM bout duration with females having longer REM bouts than males (C); furthermore, REM bout duration was significantly higher during the light period in susceptible males and females, when compared to resilient males and females (C). Side brackets indicate significant ANOVA main effect of sex or interaction p < 0.05; *, p < 0.05, Holm-Sídak’s multiple comparisons test. All plots show mean +/-SEM (both error bars and shading). n=12 females, n=8 males. Statistical details can be found in Table S1.

To observe the homeostatic regulation of sleep, we subjected mice to 6h of sleep restriction beginning at light onset and observed recovery sleep for 18h (**Figure 2B**). Female mice lost an average of 146.5 ± 4.6 min NREM (mean ± S.E.M.) sleep and 19.2 ± 2.5 min REM sleep during sleep restriction, whereas male mice lost 152.9 ± 3.6 min NREM sleep and 20.4 ± 3.0 min REM sleep (**Figure 4A, left top**). There were no differences in the amount of NREM or REM sleep lost during sleep restriction, between sexes or between behavioral phenotypes (**Figure 4A, left**). During recovery from sleep restriction, both susceptible and resilient females obtained similar amounts of NREM sleep during the 18 h recovery period (**Figure 4A, center top**). Male susceptible and resilient mice also recovered similar amounts of NREM sleep, however, there was a nonsignificant trend indicating increased sleep recovery in resilient male mice (**Figure 4A, center bottom**). Notably, the slow wave energy (SWE, the cumulative sum of SWA) that accumulated after sleep restriction was significantly increased in susceptible females when compared to resilient females (**Figure 4A, right**). A similar, but nonsignificant, increase was seen in male susceptible mice (**Figure 4A, right**). NREM SWA (a measure of sleep intensity) was also increased in the frontal and parietal EEG of susceptible females immediately after sleep restriction, relative to baseline SWA, but elevated only briefly and remained below resilient mice for the remainder of the day (**Figure 4B, left**). NREM SWA in susceptible males was increased immediately after stress in the parietal EEG and then returned to baseline for the remainder of the day (**Figure 4B right**). Resilient female and male SWA was not significantly different from baseline in either EEG lead. Together, these data indicate that SWA is elevated in resilient females (compared to susceptible females) and susceptible males (compared to resilient females) after sleep deprivation; however, this difference is not seen in SWE, an effect that may be caused by differences (nonsignificant) in sleep amount.

**Figure 4.**
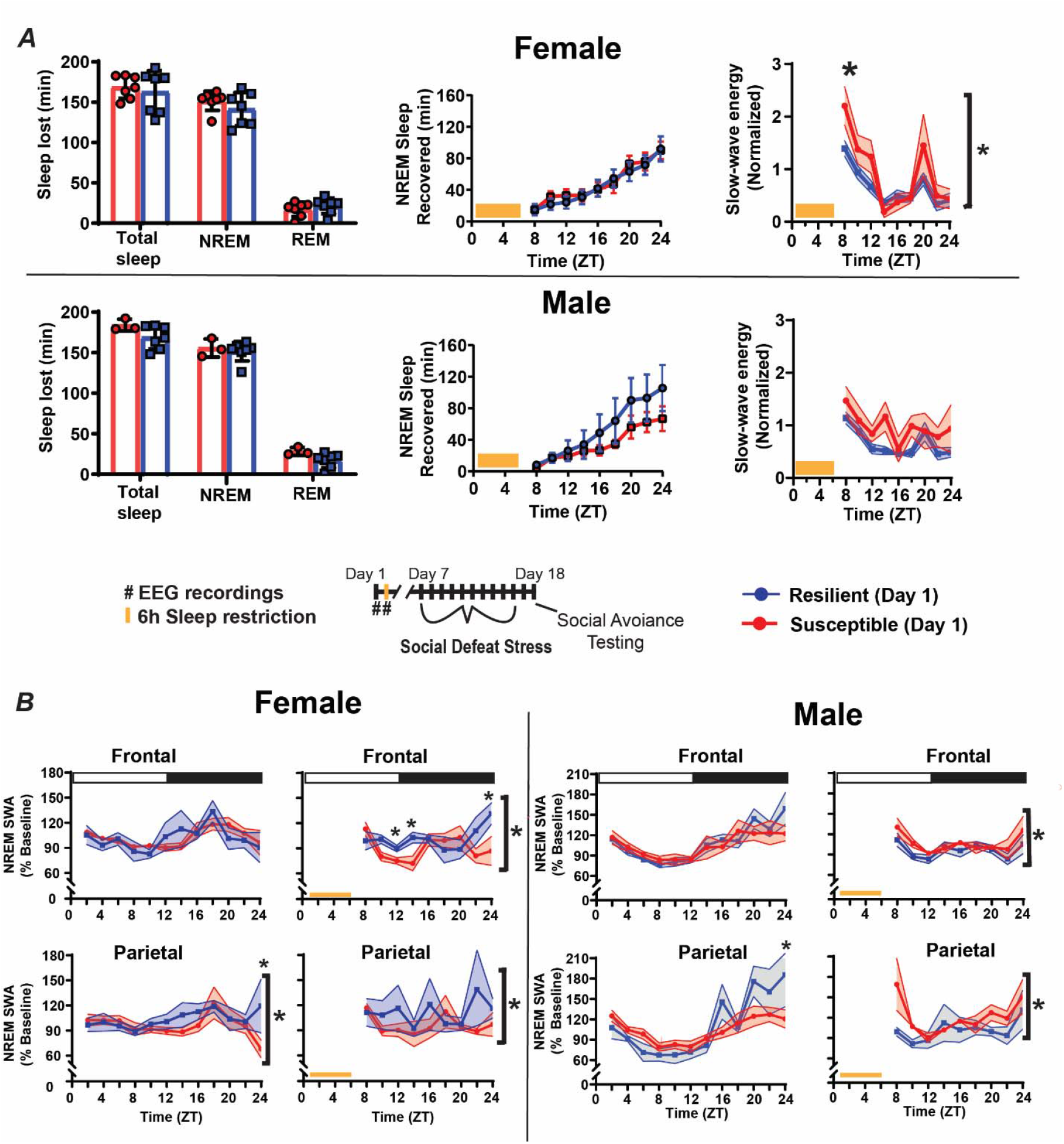
Homeostatic response to sleep deprivation before social-defeat stress: To assess differences in the homeostatic regulation of sleep, male – female pairs underwent sleep restriction during the first six hours of the light (inactive) period (A, bottom). Sleep lost during the sleep-restriction period did not differ between resilient and susceptible mice for either males or females (A, left). There were also no significant differences in amount of NREM sleep recovered between resilinet and susceptible mice, for either males or females, over the 18h recovery period (A, middle); however, there was a trend indicating resilent males may recover more NREM sleep than susceptible males. Notably, a significant increase in NREM slow-wave energy (SWE) was found in susceptible females compared to resilient females immediately after sleep restriction (A, right). This SWE increase was not seen in males (A, right). Furthermore, after sleep restriction, NREMS SWA was increased in resilient females (compared to susceptible; B, left) whereas NREM SWA was increased in susceptible males (compared to resilient; B, right). Points are mean +/-SEM (both error bars and shading). Brackets indicate significant ANOVA main effect or interaction p < 0.05; *, p < 0.05, Holm-Sídak’s multiple comparisons test. Statistical details can be found in Table S1.

Sleep amount was also significantly altered immediately after ten days of social defeat stress (**Figure 5A**). Susceptible females had significantly increased NREM sleep time when compared to baseline amounts (**Figure 5A**). This increase was caused by the duration of NREM bouts for these susceptible mice, compared to mice identified as resilient, although NREM bout duration for this comparison was not statistically significant (p = 0.063; **Figure 5A, right**). In contrast to females, NREM sleep was increased, as expected from our previously published study, after social-defeat stress in resilient males. NREM SWA was also altered near the end of the active period in both males and females (**Figure 5B**). This SWA change occurred exclusively in susceptible females and both susceptible and resilient males. For resilient males, NREM SWA decreased below baseline levels at the end of the active period, whereas, in both susceptible males and females NREM SWA fluctuated and was above or equal to baseline values during the late, active period (**Figure 5B**). The duration of REM-bouts in resilient or susceptible mice was not altered by social defeat stress (Figure 5A).

**Figure 5.**
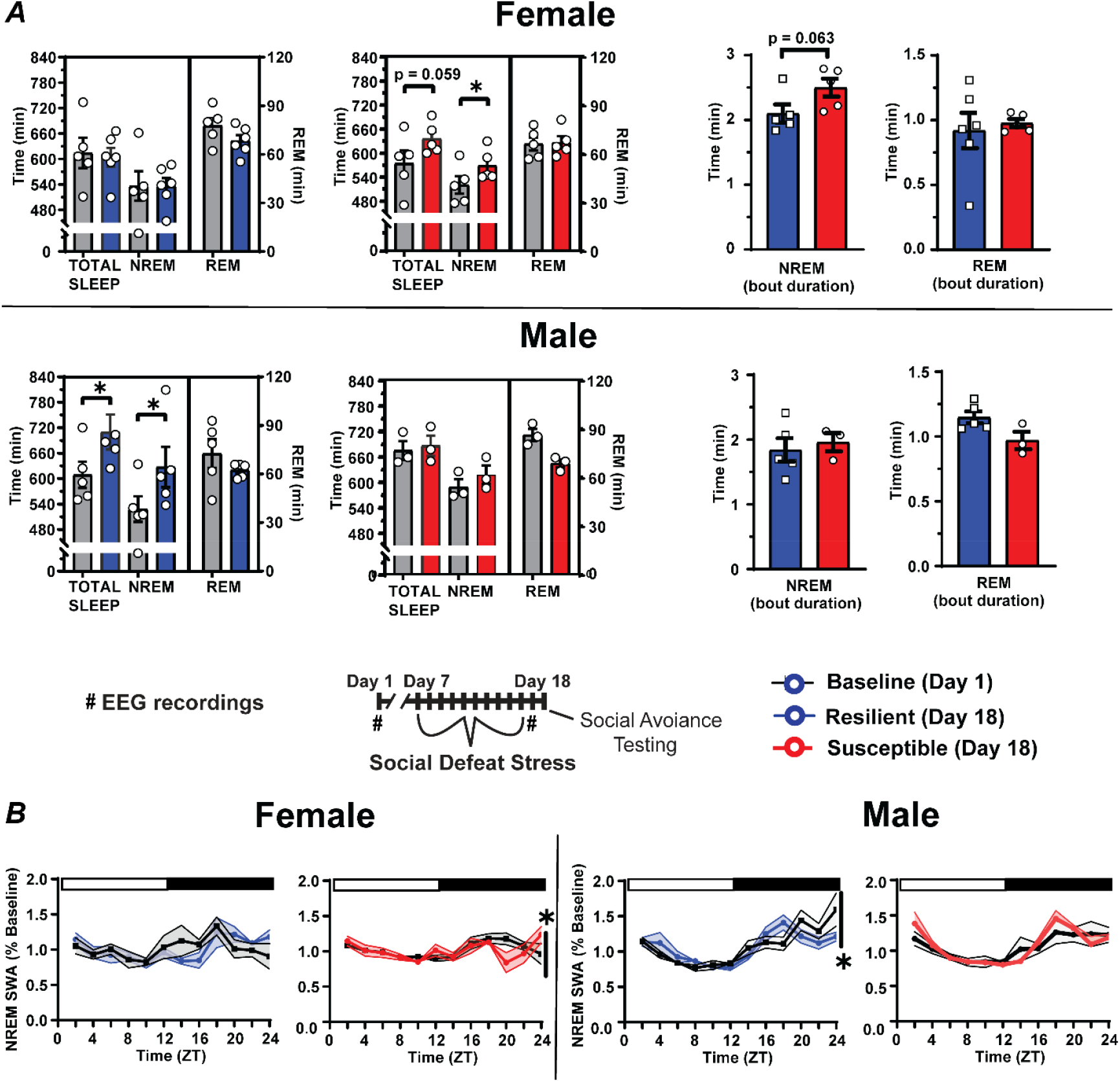
Sex differences in sleep following social-defeat stress: Female mice identified as susceptible spent significantly more time in non-rapid eye movement (NREM) sleep after social defeat stress, when compared to pre-stress sleep (A, top). A non-significant trend suggests that this was due to an increased duration of NREM bouts in susceptible female mice (A, top right). In contrast, male mice identified as resilient had increased NREM sleep time, when compared to pre-stress sleep (A, bottom). Sleep changes in susceptible females were associated with altered NREM slow-wave activity (SWA) during the active period (B left; ZT 12-24). NREM SWA in males, in contrast, was significantly altered in resilient mice during the active period (B right; ZT 12-24). Points are mean +/-SEM (both error bars and shading). Asterisks identify statistical significance. Brackets indicate significant ANOVA main effect of sex or interaction p < 0.05; *, p < 0.05. Statistical details can be found in Table S1.

## Discussion

The present study modified a recently published, non-discriminatory, social-defeat stress paradigm [23] to directly compare sleep between male and female mice. Our paradigm successfully induced stress-susceptible and stress-resilient subpopulations of both male and female mice. Fifty percent of males and sixty percent of females were classified as susceptible to social-defeat stress (**Figure 1, S1**). This distribution of susceptibility and resilience was similar to several methods of female social defeat stress reported in the literature [28, 29]. Notably, when susceptible and resilient mice were examined separately, the interaction ratios of females, but not males, change significantly from before to after social defeat stress (**Figure S1B**); however, there were similar trends in interaction ratios for both sexes and interaction time with a novel CD-1 mouse was significantly altered after stress in both sexes (**Figure S1A**). Thus, sex differences in the interaction ratios resulted from differences in male exploratory activity during the empty cage trial and not from differences in social behavior (see methods). Overall, these results indicate that the social-defeat stress paradigm, when integrated with EEG recordings, induced similar behavioral changes associated with social interaction that were not influenced by sex.

Analysis of sleep recordings both before and after social-defeat stress revealed sexually dimorphic NREM sleep patterns that are associated with resilience. One of the most striking sex differences occurred after stress, where susceptibility was associated with increased post-defeat NREM sleep exclusively in females (**Figure 5A**). This contrasts with current and past findings in males, where NREM sleep was increased only in resilient males [22]. The NREM-sleep differences we observed were due, in part, to a greater number of NREM sleep bouts in susceptible females, although this difference did not reach statistical significance (p= 0.063). Further sex differences were also revealed in NREM sleep intensity (SWA), with NREM SWA increasing in susceptible females (at the end of the dark period; Figure 5B) and decreasing in resilient males. Although sleep-differences predicting vulnerability to social defeat stress are reported in the male literature [19–22], to our knowledge, the present study is the first to investigate females and the first to conduct direct male – female comparisons. The contrasting NREM-sleep responses reported here may represent a fundamental sex-difference in the mechanisms governing resilience to social-defeat stress.

We previously demonstrated a determinative role for NREM sleep in resilience using a male social-defeat stress model [27]; however, our present data do not support the existence of this relationship in females. Notably, our findings suggest that the interaction of NREM sleep and resilience is drastically different between females and males; NREM-sleep loss in females, unlike males, may exacerbate maladaptive behavioral responses to stress. One potential cause of these sex differences is an underlying, sexually dimorphic, response by corticolimbic circuitry. Our prior findings suggest that NREM-related changes predicting resilience to social defeat stress are prominent in the medial prefrontal cortex (mPFC), a major component of this corticolimbic circuitry. Sex differences in the responsiveness of the mPFC to social stress are reported in rats [30–32]; thus, the mPFC may contribute to the sex differences observed in this study. It is also important to consider stress sensitivity when interpreting these sex differences. That is, higher or lower levels of stress may be required to elicit the same response in females as males. Importantly, females experienced fewer aggressive interactions during social defeat stress compared to males (Figure S1). Thus, this difference in stress exposure could differentially impact sleep. Although this difference in stress exposure or responsiveness may have influenced our results, the data suggest that other factors are also involved. First, sex differences in the behavioral response to stress were nearly equal. There were similar numbers of resilient mice (Figure 1) and similar changes in behavior after stress between males and females, despite a difference in aggressive interactions (Figure S1). Second, sleep was changed by stress in both males and females, suggesting that the central nervous systems of both sexes responded to stress. Lastly, sex differences in sleep were observed in resilient and susceptible populations before any exposure to stress. These pre-existing differences, especially in the homeostatic regulation of sleep, indicate that sex-differences in sleep regulation are pre-existing and, therefore, stress independent. Thus, although we cannot rule out the contribution of stress responsivity, our findings strongly support a fundamental sex difference in sleep-responses that are dependent on the resilience phenotype. This underscores the importance of understanding this interaction in females and for future studies to investigate the causal role of NREM sleep in female stress responses to test these hypotheses.

Sex-differences in sleep, before social-defeat stress, were dependent on the resilience phenotype. (**Figure 2**). These pre-existing sex differences occurred in the NREM sleep of mice resilient to, and REM sleep of mice susceptible to the effects of social-defeat stress. Overall sex-differences in sleep intensity during the active period (dark period), measured by NREM slow wave activity (**Figure 2A, top right**), were the result of differences occurring exclusively in resilient mice (**Figure 2A, middle right**). Overall differences in REM sleep during the light period (**Figure 2A, bottom**), in contrast, resulted from REM-sleep that was altered only in susceptible mice (compared to susceptible females; **Figure 2A middle**); a change that resulted from increased REM-bout length (**Figure 3C**). To our knowledge, these REM sex-differences during the light period are not reported in the existing literature. Prior studies have reported that NREM sleep differences predict resilience [22, 23, 27], thus, the significance of these REM differences are not clear. Methodological differences (discussed later) complicate comparisons of the present study with published social defeat studies. Further investigation will be required to determine if these differences associated with REM and NREM-SWA are robust between social-defeat stress models. Nevertheless, these findings reveal a significant interaction between resilience to social-defeat stress and sex; the overall quality of NREM sleep is higher in resilient males while the amount of REM sleep is increased in susceptible males—when compared to females of the same behavioral phenotype.

We assessed changes in sleep regulation to investigate the origins of sleep differences observed before and after stress. Male – female pairs were subjected to six hours of sleep restriction to monitor the recovery response to sleep loss. This is a standard paradigm that is used to quantify the homeostatic process regulating sleep. After sleep restriction, NREM slow-wave energy (SWE) returned to baseline earlier in resilient females when compared to susceptible mice (**Figure 4A top**). Resilient females also had elevated NREM sleep intensity (SWA) during recovery from sleep restriction (**Figure 4B, left)**, a difference that did not occur in males (**Figure 4B, right**). Together, these results identify a sex difference in the homeostatic mechanism regulating NREM sleep as a potential cause of the sleep differences we observed, both before and after stress. Importantly, the data also indicate that the underlying cause of SWE changes during recovery is different between sexes. SWE measures the accumulation of SWA during NREM sleep (energy = NREM SWA x NREM time). Thus, SWE can accumulate through increased sleep intensity (NREM SWA) and/or increased NREM sleep duration. Our findings suggest that resilient females (**Figure 4B, left**), but not males (**Figure 4B, right**), recover SWE from elevated NREM intensity (SWA). Resilient males, in contrast, displayed a non-significant trend indicating that NREM-sleep duration during recovery was greater than susceptible males (**Figure 4A, bottom**), thus suggesting males recover SWE by increasing NREM sleep duration. Together, these findings show that sleep recovery in females was driven by NREM-sleep intensity (NREM SWA), whereas sleep recovery males was driven by NREM-sleep duration. This difference was also associated with an increase in the contribution of SWA to the overall EEG (relative SWA) of resilient males (compared to susceptible males, **Figure 3B**). In addition, the number of NREM bouts was significantly higher in males when compared to females (**Figure 3C**). Our findings suggest some fundamental differences in sleep regulation that are dependent on the interaction of stress-resilience and sex. These differences in the homeostatic process regulating sleep appear to involve different mechanisms of recovery, NREM intensity (SWA) in females and NREM sleep duration in males.

Notably, the only other male – female mouse comparisons we are aware of in the literature [33] also reported increased NREM SWA in males (compared to females). Increased NREM amount in males was also reported during the active phase and our results did not replicate this change in NREM amount; however, there are important methodological differences that prevent direct comparisons. Mice in other studies were singly housed during sleep recording, whereas our mice were housed as male – female pairs. Thus, mice in our study had continuous sensory contact and social interaction (separated by a perforated barrier) and this is likely to alter sleep. Another potential source of variability in these studies is the female estrous cycle. Although sleep changes, especially in the active period, are reported across the estrous cycle of rats [34, 35], this does not seem to occur in C57BL/6J mice. Thus, the estrous cycle is not likely to have a major impact on our findings.

A consistent finding across this study and the existing literature is that increased NREM sleep and NREM SWA are associated with resilience in males [24, 27]. Notably, we previously demonstrated that increased NREM sleep on each day of social-defeat stress induces a resilient phenotype [27]. In the present study, we identified a sex difference in sleep-responses to stress and pre-existing sex-differences in the homeostatic mechanism driving this sleep response. Although the present study did not address causation, our findings further strengthen the importance of NREM SWA in male resilience. Importantly, they also suggest that this relationship does not exist in females. The exact mechanism involved in these effects of sleep on resilience to social-defeat stress, however, remains unknown. Slow-wave activity is the defining feature of NREM sleep and is the result of synchronous activity among cortical cells. Thus, it is reasonable to propose that the effects of NREM SWA on resilience involves the cortex. A large body of literature also demonstrates the importance of the medial prefrontal cortex (mPFC) in regulating resilience to social defeat stress [36–43]. For these reasons, it will be important for future studies to investigate how NREM SWA alters the activity of these mPFC neurons.

## Author Contributions

All authors made substantial contributions to the conception/design (BJB, ZQ, EA, JCE), data acquisition (AM, BJB, HJ, GC, EA, AF, AA, ZQ), data analysis (AM, BJB, HJ, GC, EA, AF, AA, ZQ, JCE), data interpretation (BJB, ZQ, EA, AA, KNP, JCE) or drafting/editing of the manuscript (BJB, EA, AF, AA, KNP JCE).

## Data Availability

Datasets are available upon request.

