## Supplementary material for "Sleep patterns predicting stress resilience are dependent on sex": Figure S1 and Table S1

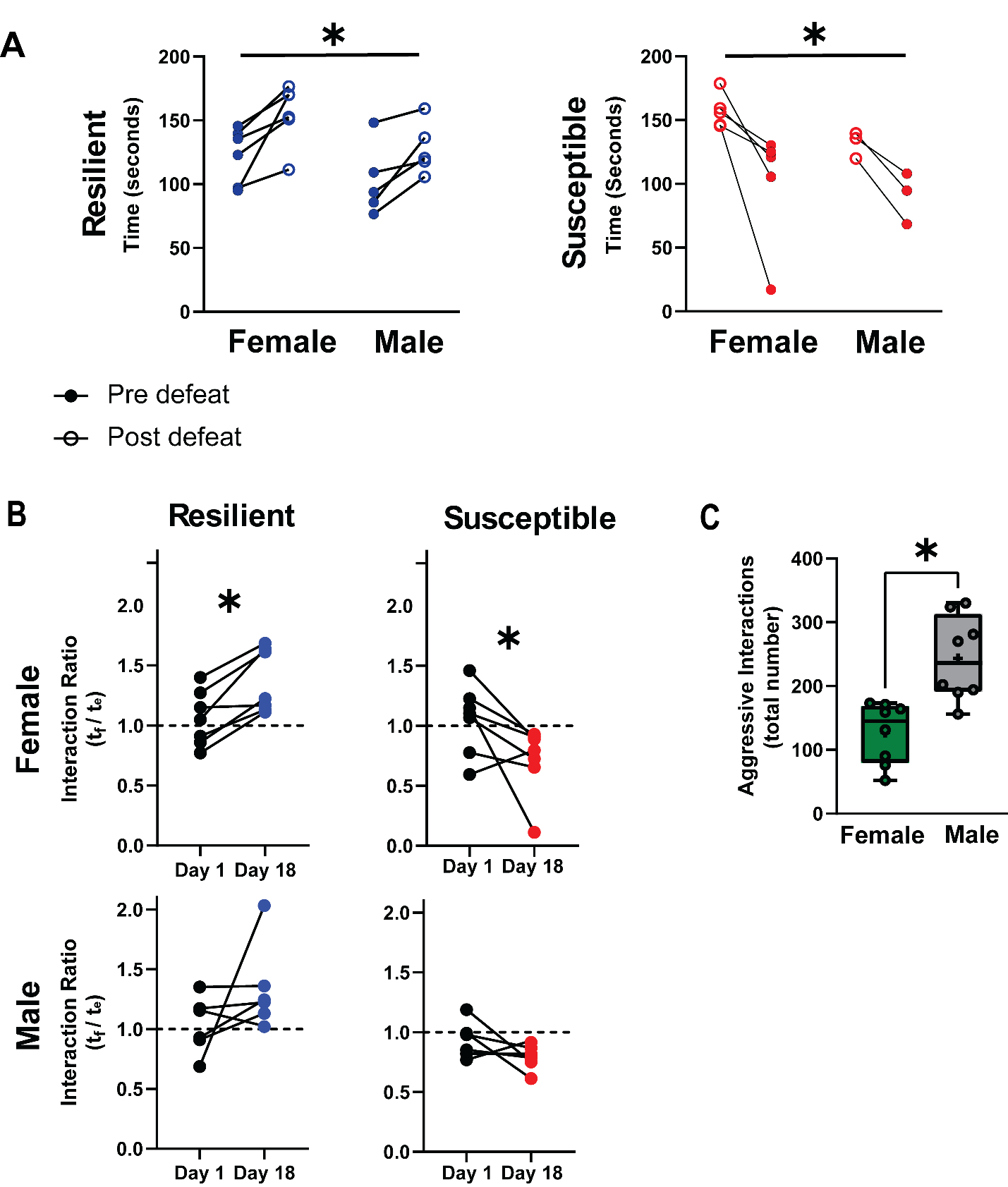
**Figure S1. Social interaction testing and social defeat stress:** Data from Figure 1D is presented by resilience phenotype to better illustrate trends (A & B). Time interacting with a caged, novel, CD1 mouse both before and after ten days of social defeat stress. Animals identified as susceptible or resilient after social defeat stress are shown separately (A). Social interaction ratios were calculated as the interaction time in A divided by time in this same region without a CD1-mouse present. Interaction scores greater than 1.1 indicated resilience, less than 0.9 indicated susceptibility (B). Box plot shows the total number of aggressive encounters experienced by males and females over the ten-day social defeat stress paradigm (C; interquartile range, median and min-max). n=10 males, n=14 females. Horizontal lines indicate significant two-way ANOVA main effect of time, p < 0.05; *, p < 0.05 paired Student’s t, Statistical details can be found in Table S1.

| **Figure** | **Independent**  **variable** | **Test** | **Measures** | ***df*** | ***F or t*** | ***P*** |
| --- | --- | --- | --- | --- | --- | --- |
| **Figure 1** | Distance traveled empty pre | Unpaired Student’s t |  | 20 | 0.029 | 0.9769 |
|  | Distance traveled empty post | Unpaired Student’s t |  | 20 | 0.781 | 0.4476 |
|  | Distance traveled full pre | Unpaired Student’s t |  | 20 | 0.605 | 0.5554 |
|  | Distance traveled full post | Unpaired Student’s t |  | 20 | 0.381 | 0.7092 |
| **Figure 2** | REM time | Two-way ANOVA | M.E. of sex | 1, 189 | 0.806 | 0.3798 |
|  |  |  | M.E. of time | 11, 189 | 23.45 | **<0.0001** |
|  |  |  | Interaction | 11, 189 | 1.558 | 0.1050 |
|  | NREM time | Two-way ANOVA | M.E. of sex | 1, 189 | 5.479 | **0.0297** |
|  |  |  | M.E. of time | 11,189 | 26.93 | **<0.0001** |
|  |  |  | Interaction | 11,189 | 1.012 | 0.4367 |
|  | SWA | Two-way ANOVA | M.E. of sex | 1, 189 | 0.004 | 0.9432 |
|  |  |  | M.E. of time | 11, 189 | 5.343 | **<0.0001** |
|  |  |  | Interaction | 11, 189 | 3.328 | **0.0003** |
| **Figure 2 & 3** | REM time | Three-way ANOVA | M.E. of sex | 1, 240 | 14.41 | **0.0003** |
|  |  |  | M.E. of resilience | 1, 240 | 0.430 | 0.5125 |
|  |  |  | M.E. of time | 11, 240 | 20.08 | **<0.0001** |
|  |  |  | Interaction of sex and resilience | 1, 240 | 3.550 | 0.0636 |
|  |  |  | Interaction of sex and time | 11, 240 | 1.565 | 0.1278 |
|  |  |  | Interaction of resilience and time | 11. 240 | 0.887 | 0.5542 |
|  |  |  | Three-way Interaction | 11, 240 | 1.140 | 0.3442 |
|  | NREM time | Three-way ANOVA | M.E. of sex | 1, 240 | 0.227 | 0.6347 |
|  |  |  | M.E. of resilience | 1, 240 | 0.063 | 0.8141 |
|  |  |  | M.E. of time | 11, 240 | 21.26 | **<0.0001** |
|  |  |  | Interaction of sex and resilience | 1, 240 | 0.127 | 0.7226 |
|  |  |  | Interaction of sex and time | 11, 240 | 0.805 | 0.6340 |
|  |  |  | Interaction of resilience and time | 11. 240 | 0.573 | 0.8481 |
|  |  |  | Three-way Interaction | 11, 240 | 0.662 | 0.7688 |
|  | SWA time | Three-way ANOVA | M.E. of sex | 1, 240 | 0.003 | 0.9551 |
|  |  |  | M.E. of resilience | 1, 240 | 595.4 | **<0.0001** |
|  |  |  | M.E. of time | 11, 240 | 16.02 | **<0.0001** |
|  |  |  | Interaction of sex and resilience | 1, 240 | 1.813 | 0.1824 |
|  |  |  | Interaction of sex and time | 11, 240 | 0.461 | 0.9208 |
|  |  |  | Interaction of resilience and time | 11. 240 | 7.780 | **<0.0001** |
|  |  |  | Three-way Interaction | 11, 240 | 0.941 | 0.4230 |
| **Figure 3** | NREM bout duration Total | Two-way ANOVA | M.E. of sex | 1,18 | 0.078 | 0.7827 |
|  |  |  | M.E. of resilience | 1,18 | 0.893 | 0.3579 |
|  |  |  | Interaction | 1,18 | 0.057 | 0.8131 |
|  | NREM bout no. Total | Two-way ANOVA | M.E. of sex | 1,18 | 3.799 | 0.0680 |
|  |  |  | M.E. of resilience | 1,18 | 0.165 | 0.6892 |
|  |  |  | Interaction | 1,18 | 1.122 | 0.3043 |
|  | REM bout duration Total | Two-way ANOVA | M.E. of sex | 1,18 | 4.223 | **0.0035** |
|  |  |  | M.E. of resilience | 1,18 | 2.512 | 0.0556 |
|  |  |  | Interaction | 1,18 | 0.842 | 0.3716 |
|  | REM bout no. Total | Two-way ANOVA | M.E. of sex | 1,18 | 2.976 | 0.1027 |
|  |  |  | M.E. of resilience | 1,18 | 0.001 | 0.9699 |
|  |  |  | Interaction | 1,18 | 2.976 | 0.1027 |
|  | NREM bout duration Light | Two-way ANOVA | M.E. of sex | 1,18 | 0.3343 | 0.5707 |
|  |  |  | M.E. of resilience | 1,18 | 1.028 | 0.3248 |
|  |  |  | Interaction | 1,18 | 0.1886 | 0.6696 |
|  | NREM bout no. Light | Two-way ANOVA | M.E. of sex | 1,18 | 1.068 | 0.3158 |
|  |  |  | M.E. of resilience | 1,18 | 1.403 | 0.2525 |
|  |  |  | Interaction | 1,18 | 0.3060 | 0.4324 |
|  | REM bout duration Light | Two-way ANOVA | M.E. of sex | 1,18 | 1.027 | 0.3250 |
|  |  |  | M.E. of resilience | 1,18 | 2.431 | 0.1374 |
|  |  |  | Interaction | 1,18 | 4.523 | **0.0484** |
|  | REM bout no. Light | Two-way ANOVA | M.E. of sex | 1,18 | 2.580 | 0.1266 |
|  |  |  | M.E. of resilience | 1,18 | 2.426 | 0.1377 |
|  |  |  | Interaction | 1,18 | 0.1333 | 0.7196 |
|  | NREM bout duration Dark | Two-way ANOVA | M.E. of sex | 1,18 | 0.6870 | 0.4187 |
|  |  |  | M.E. of resilience | 1,18 | 0.2658 | 0.6128 |
|  |  |  | Interaction | 1,18 | 0.017 | 0.8963 |
|  | NREM bout no. Dark | Two-way ANOVA | M.E. of sex | 1,18 | 5.239 | **0.0352** |
|  |  |  | M.E. of resilience | 1,18 | 0.9871 | 0.3344 |
|  |  |  | Interaction | 1,18 | 1.490 | 0.2389 |
|  | REM bout duration Dark | Two-way ANOVA | M.E. of sex | 1,18 | 0.410 | 0.5304 |
|  |  |  | M.E. of resilience | 1,18 | 1.534 | 0.2324 |
|  |  |  | Interaction | 1,18 | 0.070 | 0.7936 |
|  | REM bout no. Dark | Two-way ANOVA | M.E. of sex | 1,18 | 1.539 | 0.2316 |
|  |  |  | M.E. of resilience | 1,18 | 1.985 | 0.1769 |
|  |  |  | Interaction | 1,18 | 1.150 | 0.2986 |
| **Figure 4** | Sleep lost male Total | Unpaired Student’s t |  | 7 | 1.708 | 0.1260 |
|  | Sleep lost male NREM | Unpaired Student’s t |  | 7 | 0.491 | 0.6365 |
|  | Sleep lost male REM | Unpaired Student’s t |  | 7 | 1.965 | 0.0850 |
|  | Sleep lost female Total | Unpaired Student’s t |  | 11 | 0.541 | 0.5987 |
|  | Sleep lost female NREM | Unpaired Student’s t |  | 11 | 1.152 | 0.2718 |
|  | Sleep lost female REM | Unpaired Student’s t |  | 11 | 0.850 | 0.4117 |
|  | NREM Sleep Recovered Male | Two-way ANOVA | M.E. of Resilience | 1, 56 | 0.7120 | 0.4267 |
|  |  |  | M.E. of Time | 8, 56 | 16.350 | **<0.0001** |
|  |  |  | Interaction | 8, 56 | 1.074 | 0.3945 |
|  | NREM Sleep Recovered Female | Two-way ANOVA | M.E. of Resilience | 1, 88 | 0.045 | 0.8350 |
|  |  |  | M.E. of Time | 8, 88 | 62.13 | **<0.0001** |
|  |  |  | Interaction | 8, 88 | 0.837 | 0.5721 |
|  | Slow wave energy male | Two-way ANOVA | M.E. of Resilience | 1, 56 | 2.998 | 0.1341 |
|  |  |  | M.E. of Time | 8, 56 | 4.566 | **0.0006** |
|  |  |  | Interaction | 8, 56 | 0.803 | 0.6036 |
|  | Slow wave energy female | Two-way ANOVA | M.E. of Resilience | 1, 88 | 2.672 | 0.1262 |
|  |  |  | M.E. of Time | 8, 88 | 23.82 | **<0.0001** |
|  |  |  | Interaction | 8, 88 | 2.840 | **0.0136** |
|  | SWA baseline, frontal, male | Two-way ANOVA | M.E. of Resilience | 1, 52 | 1.020 | 0.3421 |
|  |  |  | M.E. of Time | 11, 52 | 7.386 | **<0.0001** |
|  |  |  | Interaction | 11, 52 | 1.476 | 0.2972 |
|  | SWA baseline, parietal, male | Two-way ANOVA | M.E. of Resilience | 1, 40 | 0.515 | 0.4766 |
|  |  |  | M.E. of Time | 8, 40 | 6.377 | **<0.0001** |
|  |  |  | Interaction | 8, 40 | 1.729 | 0.1170 |
|  | SWA sleep, frontal, male | Two-way ANOVA | M.E. of Resilience | 1, 52 | 7.822 | **0.0208** |
|  |  |  | M.E. of Time | 11, 52 | 1.593 | 0.1570 |
|  |  |  | Interaction | 11, 52 | 0.321 | 0.9377 |
|  | SWA sleep, parietal, male | Two-way ANOVA | M.E. of Resilience | 1, 40 | 13.72 | **0.0049** |
|  |  |  | M.E. of Time | 8, 40 | 0.9918 | 0.4567 |
|  |  |  | Interaction | 8, 40 | 0.6990 | 0.6877 |
|  | SWA baseline, frontal, female | Two-way ANOVA | M.E. of Resilience | 1, 82 | 0.0005 | 0.9812 |
|  |  |  | M.E. of Time | 11, 82 | 2.827 | **0.0082** |
|  |  |  | Interaction | 11, 82 | 0.864 | 0.5497 |
|  | SWA bsln., parietal, female | Two-way ANOVA | M.E. of Resilience | 1, 50 | 1.139 | **0.0423** |
|  |  |  | M.E. of Time | 11, 50 | 4.245 | 0.3453 |
|  |  |  | Interaction | 11, 50 | 0.9227 | 0.5019 |
|  | SWA sleep, frontal, female | Two-way ANOVA | M.E. of Resilience | 1, 82 | 6.535 | **0.0177** |
|  |  |  | M.E. of Time | 11, 82 | 1.363 | 0.2358 |
|  |  |  | Interaction | 11, 82 | 2.613 | **0.0341** |
|  | SWA sleep, parietal, female | Two-way ANOVA | M.E. of Resilience | 1, 50 | 4.392 | **0.0392** |
|  |  |  | M.E. of Time | 11, 50 | 0.4766 | 0.8694 |
|  |  |  | Interaction | 11, 50 | 0.7231 | 0.6706 |
| **Figure 5** | Male resilient total sleep | Unpaired Student’s t |  | 4 | 4.420 | **0.0115** |
|  | Male resilient NREM | Unpaired Student’s t |  | 4 | 3.384 | **0.0277** |
|  | Male resilient REM | Unpaired Student’s t |  | 4 | 1.033 | 0.3599 |
|  | Male susceptible total sleep | Unpaired Student’s t |  | 2 | 0.297 | 0.7939 |
|  | Male susceptible NREM | Unpaired Student’s t |  | 2 | 0.879 | 0.4717 |
|  | Male susceptible REM | Unpaired Student’s t |  | 2 | 3.643 | 0.0678 |
|  | Female resilient total sleep | Unpaired Student’s t |  | 5 | 0.321 | 0.7550 |
|  | Female resilient NREM | Unpaired Student’s t |  | 5 | 0.348 | 0.7418 |
|  | Female resilient REM | Unpaired Student’s t |  | 5 | 0.1578 | 0.8809 |
|  | Female suscept. total sleep | Unpaired Student’s t |  | 6 | 2.611 | 0.0593 |
|  | Female susceptible NREM | Unpaired Student’s t |  | 6 | 4.362 | **0.0121** |
|  | Female susceptible REM | Unpaired Student’s t |  | 6 | 0.141 | 0.8945 |
|  | Male NREM bout duration | Unpaired Student’s t |  | 6 | 0.459 | 0.6623 |
|  | Male REM bout duration | Unpaired Student’s t |  | 6 | 2.268 | 0.0638 |
|  | Female NREM bout duration | Unpaired Student’s t |  | 11 | 2.356 | **0.0429** |
|  | Female REM bout duration | Unpaired Student’s t |  | 11 | 0.378 | 0.7139 |
|  | Male SWA resilient | Two-way ANOVA | M.E. of Resilience | 1, 90 | 0.269 | 0.6052 |
|  |  |  | M.E. of Time | 11, 90 | 11.69 | **0.0001** |
|  |  |  | Interaction | 11, 90 | 2.071 | **0.0305** |
|  | Male SWA Susc. | Two-way ANOVA | M.E. of Resilience | 1, 44 | 0.256 | 0.6383 |
|  |  |  | M.E. of Time | 11, 44 | 8.491 | **0.0091** |
|  |  |  | Interaction | 11, 44 | 0.801 | 0.6396 |
|  | Female SWA resil. | Two-way ANOVA | M.E. of Resilience | 1, 56 | 0.906 | 0.7542 |
|  |  |  | M.E. of Time | 11, 56 | 0.677 | 0.3453 |
|  |  |  | Interaction | 11, 56 | 0.717 | 0.7173 |
|  | Female SWA Sus. | Two-way ANOVA | M.E. of Resilience | 1, 176 | 3.271 | **0.0004** |
|  |  |  | M.E. of Time | 11,176 | 0.215 | 0.8066 |
|  |  |  | Interaction | 11, 176 | 1.005 | 0.4607 |
| **Figure S1** | Susceptible Mice | Two-way ANOVA | M.E. Defeat | 1, 6 | 13.04 | **0.0112** |
|  |  |  | M.E. Sex | 1, 6 | 1.053 | 0.3445 |
|  |  |  | Interaction | 1, 6 | 0.344 | 0.5788 |
|  | Resilient Mice | Two-way ANOVA | M.E. Defeat | 1, 6 | 22.53 | **0.0017** |
|  |  |  | M.E. Sex | 1, 6 | 3.330 | 0.1013 |
|  |  |  | Interaction | 1, 6 | 0.373 | 0.5566 |
|  | Aggressive interactions | Paired Student’s t |  | 7 | 6.913 | **0.0002** |
|  | Female susceptible | Paired Student’s t |  | 7 | 2.301 | **0.0493** |
|  | Female resilient | Paired Student’s t |  | 7 | 4.731 | **0.0003** |
|  | Male susceptible | Paired Student’s t |  | 6 | 1.283 | 0.2558 |
|  | Male resilient | Paired Student’s t |  | 6 | 1.446 | 0.2078 |

**Table S1. Summary of Statistical results:** Statistical tests used in each figure are shown with associated degrees of freedom, test statistics and p values. M.E., ANOVA main effect; interaction, ANOVA interaction effect; Bsln., baseline; no. number.
